# Auxin coordinates cell states during Arabidopsis root development

**DOI:** 10.64898/2026.01.02.697414

**Authors:** Cassandra Maranas, Sydney VanGilder, Linda Nguyen, Jennifer Nemhauser

## Abstract

Cell-to-cell variation in gene expression can be highly detrimental and, in some contexts, is actively buffered out; however, in other contexts, it is crucial and actively amplified. For example, variation must be minimized to build organs with consistent size and shape, yet the initiation of organogenesis requires a subset of cells to take on a new fate, a process that often relies on small differences between cells. In plants, much of development is controlled by the hormone auxin, which has been hypothesized to coordinate cell responses by inducing degradation of transcriptional repressors. To quantify the level of cell-to-cell variation and directly test its connection to auxin signaling, we assessed variation in expression of a lateral root founder cell marker *GATA23* when auxin levels or responsiveness was modulated. We found that auxin acted as both an amplifier and a constrainer of transcriptional variation during the initiation of a new root. We then extended this work to analysis of root regeneration, where auxin was also found to play a critical role in coordinating cells during fate transitions.

**ARTICLE SUMMARY:** During organogenesis, cell-to-cell variation is induced, enabling some cells to adopt a new identity and differentiate. Despite this dependence on variation, organogenesis is robust in developmental stages and outcomes. In plants, organogenesis is usually controlled by the hormone auxin, and we hypothesized that auxin ensures coordination among differentiating cells to enable robust development. Using Arabidopsis lateral root development as a model, our results showed that cells receiving the highest auxin dosage both amplify their own auxin response and repress the auxin response in neighboring cells receiving lower auxin dosage, establishing separation between differentiating and non-differentiating cells and ensuring coordinated organogenesis.

## INTRODUCTION

Stochastic cell-to-cell variation in gene expression is well established across organisms and cell types^1^. For many genes, cell-to-cell variation in expression is deleterious to functioning, and cellular mechanisms to buffer it are employed on the gene, transcript, and protein levels^2–6^. However, there is also evidence that expression variability can be advantageous, perhaps even under positive selection^7^, and reinforced by multiple regulatory mechanisms^8^. The importance of cell-to-cell variation has been established for many cell differentiation processes (including photoreceptor fate specification in retinal development^9^, blastocyst organization^10^, and embryonic pancreatic organogenesis^11^). These processes often rely on inherent transcriptional noise to trigger cell fate transitions in a subset of cells^12^. In fact, existing cell-to-cell variation can be amplified during organogenesis to form defined boundaries between the identities of differentiating and non-differentiating cells and ensure coordinated development^2^. Studies using single cell RNA sequencing have identified many genes associated with cell differentiation as being variably expressed^13,14^.

In studies of cell differentiation trajectories, one idea that has been explored is that of a “transition cell state” in which a cell’s gene expression profile bears similarities to both the undifferentiated and differentiated cell states and is more heterogeneous than either state^15,16^. Because of limitations in current experimental techniques for assessing transient cell states, an alternative approach is stochastic modeling of gene expression networks driving differentiation. One such study^17^ developed a model of cell differentiation and transition state dynamics based on Waddington’s epigenetic landscape^18^ wherein cell states were represented as energy potentials, together forming a probabilistic developmental landscape, and cell differentiation was a cell’s trajectory through this landscape. Changing the concentration of a signaling molecule distorted the energy landscape of the transition state and thus also distorted cell differentiation trajectories through it. Other work has also linked extracellular signaling to heterogeneous transition state dynamics and cell differentiation trajectories^19,20^ but the mechanisms describing how cells variably process these signals to make cell fate decisions are not yet clear.

Despite the established variability in cell states and differentiation trajectories, development is mostly predictable and robust, resulting in consistently formed tissues and organs. *Arabidopsis* sepal development has been a particularly useful model for studying the relationship between cell-to-cell heterogeneity and developmental robustness. First, in a landmark study, it was shown that when more cells are present (such as in a mature sepal), cell-to-cell variation in growth rates is spatiotemporally averaged, buffering these differences and enabling consistent sepal growth^4^. As such, initial cell-to-cell variability is resolved over time and space. Second, rapid and coordinated amplification of cell-to-cell variation is key to sepal initiation, but increasing the speed and sensitivity of the response comes at the expense of cell-to-cell coordination and robustness^21^. Third, if there is sufficient variation in the earliest stages of sepal development (patterning of sepal primordia sites by the plant hormone auxin), the lack of cell-to-cell coordination is propagated through developmental stages, resulting in inconsistently formed sepals^21,22^. Taken together, this work shows that proper development is predicated on the management of cell-to-cell variation, first amplifying it to initiate differentiation and later attenuating it to ensure robust progression through developmental stages.

Like sepals, lateral root (LR) development is governed by auxin signaling. Lateral roots are also an excellent model for understanding variation, as their initiation is regulated by one of the best understood plant morphogenetic processes and the roots themselves are completely dispensable in lab-grown plants^23^. LRs are initiated when a subset of xylem pole pericycle (XPP) cells receive a sufficiently high auxin signal. They then take on an LR founder cell identity and begin to divide (Fig. 1A)^24^. The number of LR founder cells contributing to each LR has been historically described as 2-3^25^; however, more recent work has revealed that it can vary from as few as 2 to as many as 11^26^.

**Figure 1:**
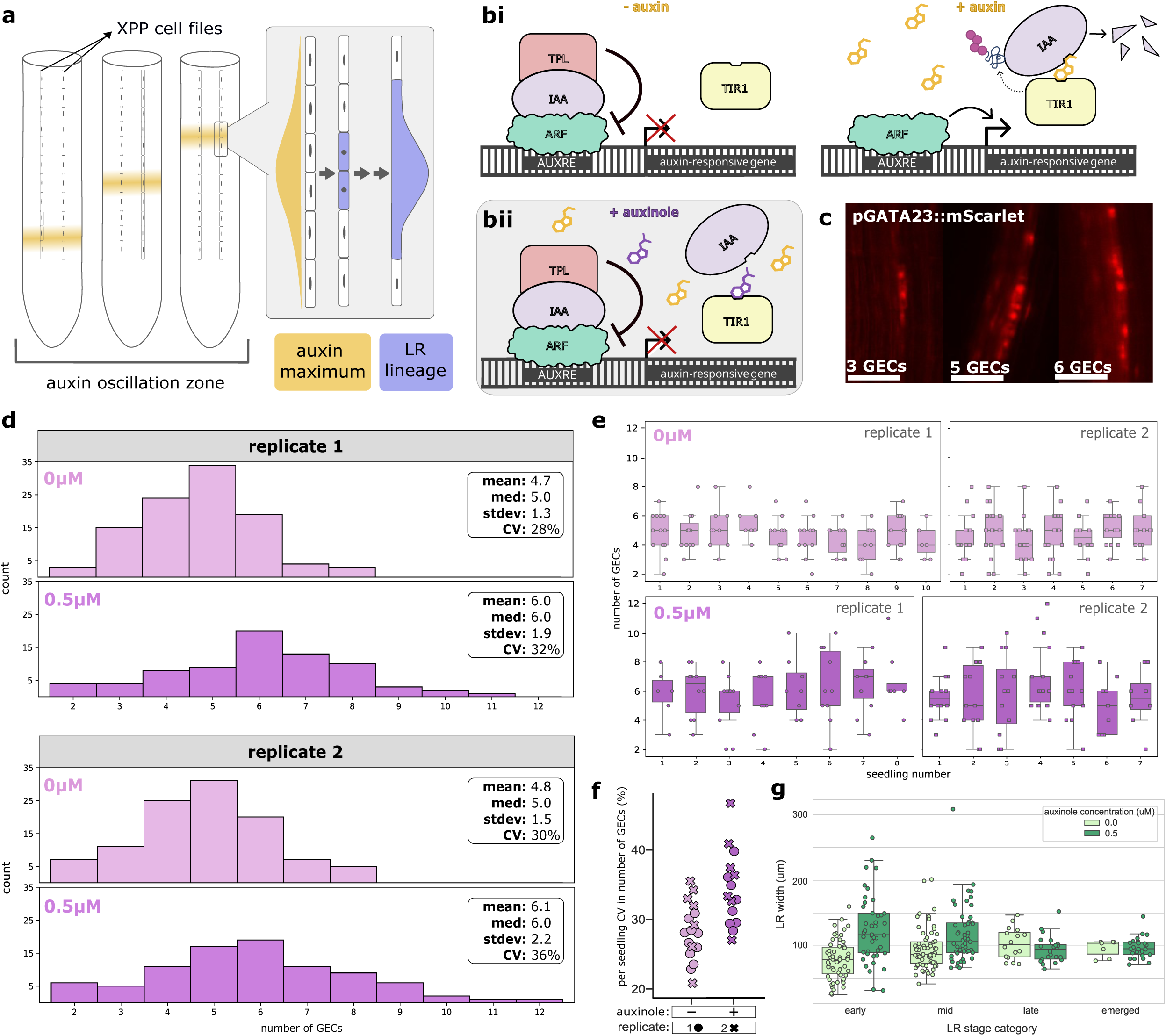
Dampening the auxin signal strength with auxinole treatment increases variability in the early stages of lateral root initiation. **A.** Specification of LR founder cells. (left) Auxin concentration oscillates along the length of the root defining LR branch sites. (right) A subset of XPP cells which are exposed to high auxin signal undergo a cell fate transition, becoming LR founder cells and dividing to form the LR. **Bi.** Auxin signaling schematics. (left) Without auxin present, TPL represses gene expression through contact with Aux/IAA. (right) When auxin is present, it mediates ubiquitination and degradation of Aux/IAA through contact with TIR1, relieving repression of auxin-responsive genes. **Bii.** With both auxin and auxinole present, auxinole competes for binding with TIR1, reducing IAA degradation rate and slowing relief of repression of auxin-responsive genes. **C.** Examples of counting GECs using the pGATA23::mScarlet reporter line. The GEC data is used to produce panels D-F. **D.** Overall distribution in GEC number in the control (light pink) and 0.5 μM auxinole treatment (magenta) for two cell counting replicates. Summary stats are included for each distribution. **E.** Per seedling distribution in GEC number for control (top) and auxinole (bottom) treatments across two replicates. **F.** Per seedling CV in GEC number for the control and 0.5 μM auxinole treatment. **G.** Overall distribution of LR widths by developmental stage. Stage categories are as follows – early: stages 1-2; mid: stages 3-5; late: stages 6-7, according to Banda et. al 2019^23^. For early stage LRs, width was measured based on the extent of GATA23 reporter expression. For the later LR categories, width was measured based on LR protrusion from the main root.

In the absence of auxin, the expression of auxin-responsive genes is repressed by the TPL/TPR corepressors which inhibit the activity of AUXIN RESPONSE FACTOR (ARF) activators through association with coreceptors/adaptors from the AUXIN/INDOLE-3-ACETIC ACID (Aux/IAA) family (Fig. 1Bi). Auxin signaling facilitates rapid Aux/IAA degradation through formation of a complex between auxin, the Aux/IAA and auxin receptors like TRANSPORT INHIBITOR RESPONSE 1 (TIR1) (Fig. 1Bi). The rate of auxin-induced degradation controls the speed of transcriptional responses^27^, as well as the rate of LR morphogenesis^28^. We hypothesized that, beyond acting as a timer for events within a single cell, auxin might also act to coordinate the response of multiple cells undergoing parallel fate changes during development. To test this hypothesis, we used a variety of chemical and genetic tools to perturb auxin signaling and measure the impact on variation in initiation of a new root in the context of the primary root as well as during regeneration. Our results support a role for auxin as a coordinator of multi-cell behaviors during development.

## RESULTS AND DISCUSSION

As a metric for variation in LR initiation and an early step in auxin signaling, we measured the number of LR founder cells. As *GATA TRANSCRIPTION FACTOR 23* (*GATA23*) expression is one of the earliest markers of LR founder cell identity^29^, we counted the number of *GATA23-*expressing cells (GECs) in early stage LRs as a proxy for the number of founder cells (Fig. 1C). In our standard growth conditions, the overall distribution of GECs was roughly symmetrical, with a peak around 5, and a range of 2-8, consistent with previous studies^26,30,31^. The coefficient of variation (CV), a normalized expression of data spread, makes it possible to compare the extent of variation between different data sets. The CV of GECs in WT plants was 28% (Fig. 1D). We observed some seedling-to-seedling variation in the distribution of GEC number, but the median number of GECs was relatively consistent between seedlings (Fig. 1E). Thus, the observed variability in GEC number is inherent to the LR initiation process, as it is recapitulated in each seedling.

To assess the impact of dampening the auxin signal, we grew seedlings in the presence of low levels of auxinole, a compound that competes with auxin for binding with TIR1 (Fig1Bii)^32^. To find the ideal concentration for these assays, we quantified emerged lateral root density in a range of doses (Fig. S1). From this result, we selected 0.5 μM auxinole, as it was able to mildly reduce but not block lateral root development. The distribution of number of GECs in the auxinole-treated LRs was considerably wider than in the control (Fig. 1D), with the maximum increasing to 12 (1.5x the control). Auxinole treatment also increased the variability by approximately 20%, resulting in a CV of 34%. As in the control conditions, the median number of GECs for LRs within each seedling is relatively consistent (Fig. 1E). Auxinole shifted the distribution of CV across seedlings upwards, while maintaining a similar range with control conditions (Fig. 1F). From these results, we can conclude that the correct level of auxin response is required to maintain robustness in the absolute number and variation in number of LR founder cells.

We were next interested in examining the impact of auxinole treatment in later stages of LR development. To capture LRs across all stages, we measured the width of each LR on each *GATA23* reporter seedling screened. Plotting these LR widths by stage (Fig. 1G) we found that early stage LRs were the most variable in width, with variation decreasing through each of the later stages. Treatment with auxinole resulted in increased average width and variation therein for early and mid-stage LRs, but in later stages the distribution grew closer to that of the control. This trend indicates that as LR development progresses, initial cell-to-cell variation is resolved, allowing robust root formation, even when the coordinating auxin signal is dampened by auxinole treatment. This trend could be a function of the increased number of cells in later stage LRs, allowing for spatiotemporal averaging of each cell’s growth and cell cycle fluctuations, similar to what is observed in sepal development^4^.

One model to explain our results is that a weakened auxin response is ineffective in repressing LR fate in cells adjacent to the founder cells, a role that has been previously documented^33^. This is consistent with the observation that auxinole treatment significantly increased the high end of the GEC distribution (Fig. 1D). Following this logic, unrepressed neighboring cells might have lower expression of LR-initiation genes like *GATA23* than cells at the core of the initiating LR. We tested this prediction with a previously characterized integrase-based durable recorder of *GATA23* expression, a tool we call an integrase switch^34^. The integrase switch has two components: (1) the target which can switch between expression of one gene to another by inverting the direction of the promoter, and (2) a driver that directs the expression of the PhiC31 serine integrase (Fig, 2A). In the integrase switch used here, PhiC31 accumulates in cells when *GATA23* is expressed, and, past a certain accumulation threshold, the target is switched so that cells express mScarlet instead of mTurq (Fig. 2B). Crucially, unlike a transcriptional reporter, the integrase switch is a binary response: only those cells that surpass a threshold level of *GATA23* expression will switch (*GATA23-*switched cells, GSCs) (Fig. 2A), and the change in the target and associated reporter expression is permanent and durable in these cells. Because of the binary nature and imposed expression threshold of the switch, we predicted that the number of GSCs would not be affected by auxinole to the same degree as GECs, because the level of *GATA23* expression in ineffectively repressed adjacent cells would likely be below the threshold at which the integrase switch occurs.

**Figure 2:**
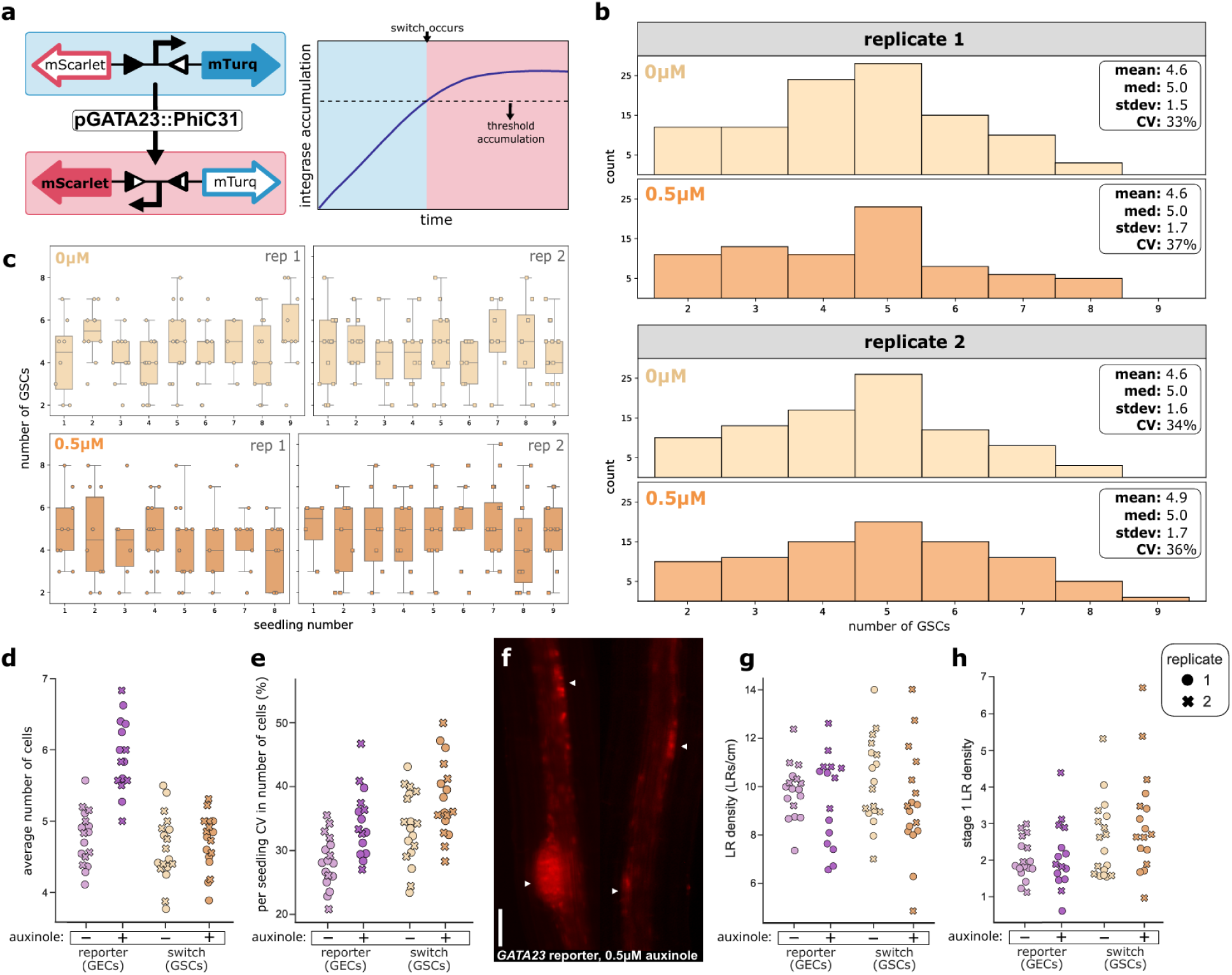
*GATA23* integrase switch shows that dampening the auxin signal during LR initiation results in incomplete repression of the auxin response in neighboring cells and an increased rate of early LR arrests. **A.** *GATA23* integrase recorder schematic. (left) The recorder starting state is expression of mTurquoise. With sufficient accumulation, the PhiC31 integrase mediates DNA recombination, inverting the promoter direction and switching expression to mScarlet. The switch is permanent, thus mScarlet expression is sustained indefinitely. **B.** Overall distribution in GSC number (as measured by counting the number of mScarlet-expressing cells in the *GATA23* switch line) in the control (light orange) and 0.5 μM auxinole treatment (dark orange) for two cell counting replicates. Summary stats are included for each distribution. **C.** Per seedling distribution in GSC number for control (top) and auxinole (bottom) treatments across two replicates. **D.** Per seedling average number of cells for the *GATA23* reporter (GECs, pink) and recorder (GSCs, orange) in the control and auxinole treatment. **E.** Per seedling CV in number of cells for the *GATA23* reporter (GECs, pink) and recorder (GSCs, orange) in the control and auxinole treatment. **F.** Example images of “double LRs” seen more frequently in auxinole-treated seedlings. Scale bar is 100 μm. **G, H.** Total (**G**) and stage 1 only (**H**) LR densities for the *GATA23* reporter and recorder, with 0 and 0.5 μM auxinole added.

Repeating the cell counting process in our *GATA23* switch line, evaluating the number of GSCs, we found a comparable distribution to that found in the transcriptional reporter. The distributions of GECs (Fig. 1C) and GSCs (Fig. 2B) had the same median of 5 cells and the same minimum (2) and maximum (8). Seedling specific plotting of the number of GSCs (Fig. 2C) also reveals similar patterns to that of GECs, with every seedling across both experiments having a median number of cells between 4 and 6 (Fig. 2C). However, unlike for the GECs, application of auxinole did not result in a significant increase in median number of GSCs. We computed the average GEC or GSC number for all counted LRs on each screened seedling in each condition and, with auxinole addition, we found a significant upwards shift in average GEC number and no significant difference in GSCs (Fig. 2D).

The variability in number of GSCs was modestly increased with auxinole treatment, but not to the same level as was found in GECs (an overall 6% increase in CV compared to 21% for GECs) (Fig. 2E). On a per seedling basis, each auxinole-treated seedling had a median number of GSCs between 4 and 6 (Fig. 2C), just like the control. As we predicted, the effect of auxinole on GSC numbers was weaker compared to GECs. Additionally, as would be expected if auxinole treatment impairs repression of the auxin response in neighboring cells, auxinole treatment led to a higher frequency of “double LRs”, where two LRs were formed in very close proximity to one another (Fig. 2F). Loss of function of the peptide RALFL34 is known to interfere with the repression of LR neighboring cells, with the *ralfl34* mutant showing increased divisions of LR-neighboring XPP cells (indicating founder cell identity) and an increased incidence of double LRs^35^. Both phenotypes are similar to our results with auxinole treatment (Fig. 2D, Fig. 2F), further supporting a model where auxin signaling levels in central GECs are optimized for lateral inhibition of auxin responsiveness in neighboring cells.

Another difference we noted with the integrase recorder was a higher relative frequency of low GSC numbers. This could be attributed to the same dampened neighbor cell repression if this phenomenon evenly affected LRs regardless of GSC number, but this is not what we observed. Another possibility is that some proportion of these low GSC LRs are LR terminations. Because many means of LR arrest occur in early stages, such as through inhibition of LR founder cell division by cytokinin^36^, we posited that the switch line should have a higher proportion of early stage LRs when compared with the transcriptional reporter, which is indeed what we observed. While the overall LR density was similar between the switch and reporter lines (Fig. 2G), Stage 1 density was higher in the switch line (Fig. 2H). Additionally, auxinole treatment increased the relative proportion of Stage 1 LRs in the recorder seedlings (Fig. S2A), but not in the switch seedlings (Fig. S2B). This suggests that auxinole treatment increases the likelihood of an LR to terminate early.

In LR development, cytokinin signaling works antagonistically to auxin signaling and it is the balance between the two that guides root development. Mutually antagonistic hormone signaling dynamics are common in plants and are known to affect variability in plant traits, such as in *Arabidopsis* seed germination, where the relative signaling strength of the hormones abscisic acid and gibberellic acid sets the level of variability in germination timing^37^. Similarly, our results suggest that shifting the balance between auxin and cytokinin signaling (in this case by reducing auxin signal strength) also affects variability in cell responses.

As an independent test of the impact of lowered auxin sensitivity on variation, we analyzed a previously characterized line expressing an IAA14 mutant with a lower binding affinity for auxin (*IAAslow*)^30^. While the overall LR density in *IAAslow* seedlings was not significantly changed compared to WT, the variation was greatly increased (Fig. 3A). The CV in number of GECs in *IAAslow* was 42%, compared to 10% in WT. With auxinole treatment, the WT CV in LR density was 15%, which increased to 47% in *IAAslow*. As reported previously, the distribution of LR stages was quite different in *IAAslow* with the majority of LRs in Stage 1 (Fig. 3B, Fig. S2C) and a dramatic reduction in emerged LRs (Fig. 3C, Fig. S2C). We predicted we would find similar increases in median and variability in number of GECs in *IAAslow* as we did with auxinole treatment (Fig. 1C). Instead, we found that the median number and range of GECs in *IAAslow* was the same as in WT (Fig. 3D,E), albeit with a slight increase in the average from 4.8 to 5.3 GECs. The range of GEC values increased in *IAAslow*, with a maximum number of GECs of 11 compared to 8 in WT.

**Figure 3:**
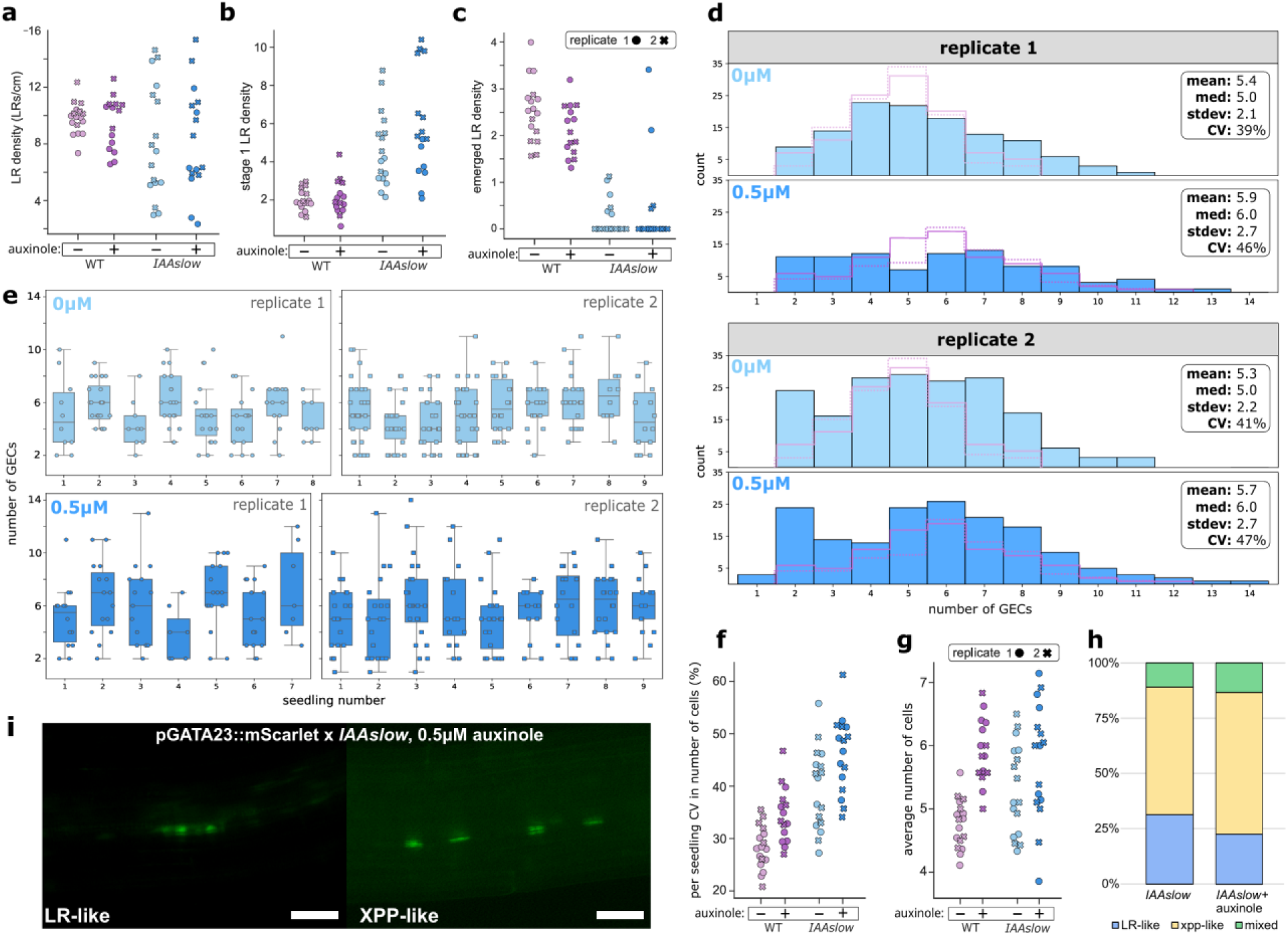
Further dampening of auxin signaling with auxinole and *IAAslow* shows differing effects on GEC number and variation therein, and potentially muddied transition state dynamics. **A-C.** Total (**A**), stage 1 only (**B**), and emerged only (**C**) LR densities in WT and *IAAslow* backgrounds, with 0 and 0.5 μM auxinole added, over two replicates. **D.** Overall distribution in GEC number in *IAAslow* in the control (light blue) and 0.5 μM auxinole treatment (dark blue) for two cell counting replicates. GECs for the *IAAslow* background were determined by counting GECs in a pGATA23::mScarlet x *IAAslow* cross line. Summary stats are included for each distribution. For reference, the 0 and 0.5 μM auxinole distributions in WT (Fig. 1D) are represented as a dotted (replicate 1) and solid (replicate 2) outlines over the *IAAslow* 0 and 0.5 μM auxinole distributions, respectively. **E.** Per seedling distribution in GEC number in *IAAslow* for control (top) and 0.5 μM auxinole (bottom) treatments across two replicates. **F, G.** Per seedling CV in (**F**) and average (**G**) GEC number in WT and *IAAslow* and with 0 and 0.5 μM auxinole treatment. **H.** Proportion of cell appearances in *IAAslow* GECs. **I.** Example images of LR-like (left) and XPP-like (right) GECs. Scale bar is 100 μm.

The *IAAslow* mutant and auxinole should act independently to reduce auxin sensitivity, so in combination might reveal the upper limits of variation. When *IAAslow* seedlings were exposed to auxinole (*IAAslow*+auxinole), the median for the number of GECs in *IAAslow*+auxinole was 6, an increase from the no auxinole control but relatively unchanged compared to WT+auxinole (Fig. 3D). Each individual seedling had a median number of GECs between 4 and 7 (Fig. 3E). The mean showed the same pattern, where the *IAAslow*+auxinole mean of 5.8 GECs was comparable to the WT+auxinole mean of 6.0 GECs. The distribution in per seedling average GEC numbers in *IAAslow*+auxinole was not significantly changed compared to *IAAslow* (Fig. 3F). This contrasts with the significant upward shift seen in WT+auxinole compared to WT alone (Fig. 3F). Looking at the variation in the number of GECs during LR initiation, we found that *IAAslow*+auxinole was the most variable, with a CV of 47%. This compares to a CV of 40% for the no auxinole *IAAslow* control and 34% for WT+auxinole. Plotting CV in GEC number on a per seedling basis for each line and in each auxinole treatment (Fig. 3G), we found that auxinole addition consistently increased the average CV, shifting the distribution upwards in both WT and *IAAslow*. So, while the combinatorial effect on variability in GEC number by *IAAslow* and auxinole was synergistic, the effect on GEC number itself was not. This pattern suggests that there is a functional limit to the number of LR initial cells (a number already close to saturation in both mutant and auxinole treatment), but perhaps not in the potential for variation.

In addition to altered number of GECs, we noticed the majority of the nuclei in GECs in the *IAAslow* background were unusually shaped, often resembling the lens-like shape of nuclei in non-LR XPP cells (Fig. 3H). These XPP-like cells made up 58% of the screened LRs and another 10% were comprised of a mix of XPP-like and more typical LR-like cells (Fig. 3I). Interestingly, the appearance of the cells in the LR had little bearing on the distribution of number of GECs. It could be that these XPP-like GECs represent a transition state within the LR initiation process. Changes to a cell’s signaling environment can alter the stability and durability of the transition state^17^. Recently, the capacity of precursor cells to undergo mixed cell fate transitions and the importance of auxin in controlling these transitions has been shown in stomatal development^38^. In *IAAslow*, auxin signal strength is much reduced, and it would make sense that a transition state could arise from a reduction in auxin signal strength in LR precursor XPP cells and/or a reduction in repression of the auxin signal strength in XPP cells neighboring LR precursors. In either case, a cell’s trajectory could be diverted towards a relatively stable transition state. If this were the case, we would expect that further dampening of the auxin signal with auxinole would increase the likelihood of cells entering this transition state. Consistent with this prediction, auxinole treatment increased the number of both XPP-like GECs to 64% (Fig. 3I).

If auxin is indeed a controller of variation during root initiation, we reasoned that similar trends should be found in regenerated roots (RRs) (Fig. 4A), which involve expression of many of the same developmental genes^39^. In root regeneration from leaf explants, the wound site is flooded with high levels of auxin which are required for RR initiation^40^. Root regeneration is well suited to analyzing phenotype variation because, compared to LR development, root regeneration is not robust. Leaf explants of the same age, and even from the same seedling, show natural variation in the number of RRs and the timing of their development^41^. In pre-emerged stages, RRs are less consistent in size and shape, while LRs have a more consistent morphology ^23^. This initial variation gets buffered, however, as mature RRs are relatively consistent in morphology^42^.

**Figure 4:**
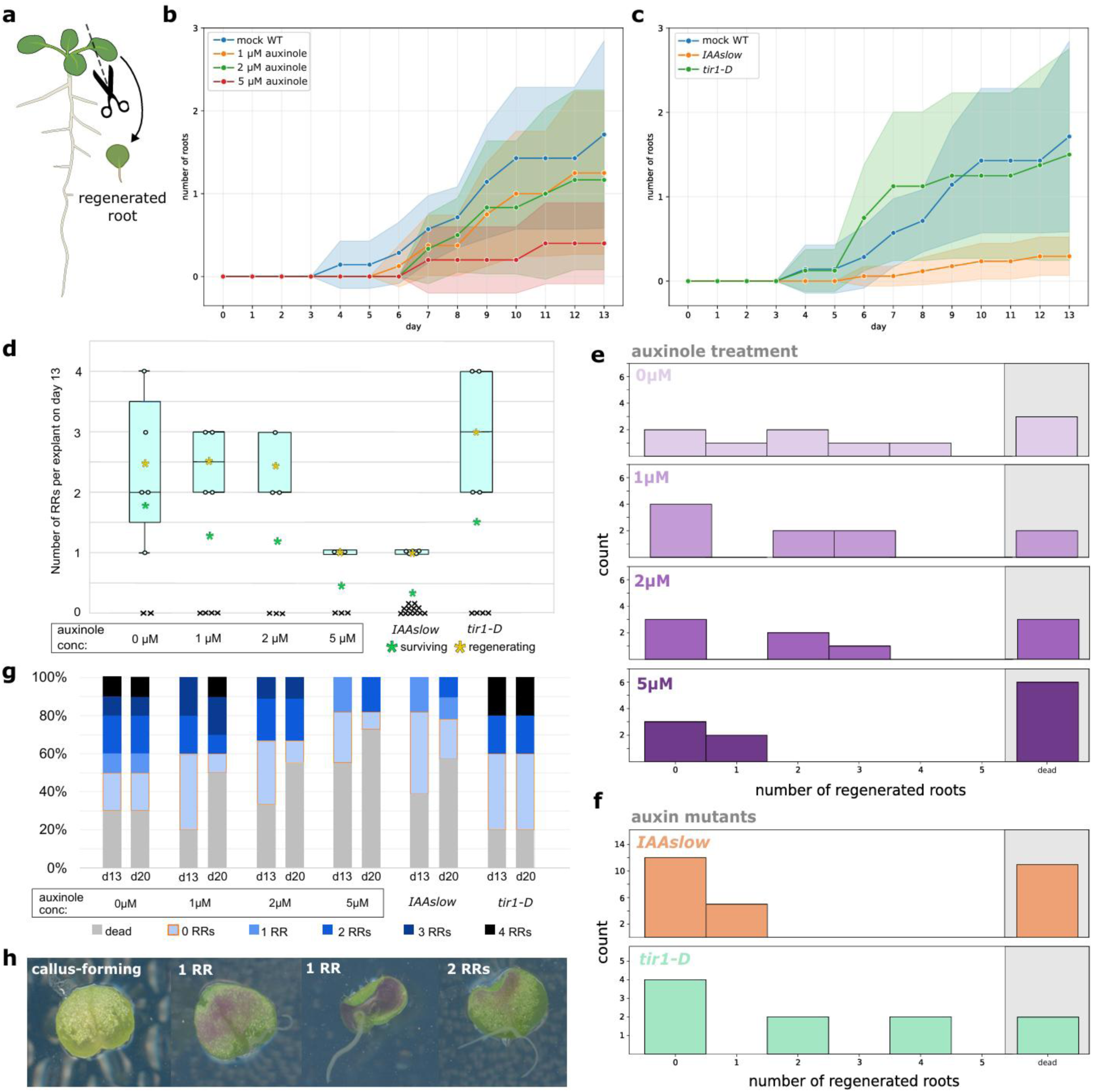
Auxin signaling conditions impact frequency and robustness of root regeneration outcomes. **A.** Schematic of root regeneration from *Arabidopsis* leaf explants. The leaf is removed from the seedling and cultured, where it may regenerate a new root/s near the wounding site. **B, C.** Regeneration timelines for (**B**) auxinole-treated and (**C**) auxin mutant explants. Day 0 is the day of explant excision. Shaded areas represent standard error. **D.** Distribution for the number of RRs on day 13 for each auxinole treatment and auxin mutant line. Each point is the number of RRs for a single explant. The green asterisks represent the average number of RRs among all surviving explants in each group, and the yellow asterisks represent the average when including only explants with at least one RR at day 13. The x data points represent surviving explants which regenerated no roots. **E, F.** Overall distribution in number of RRs for the auxinole treatments (**E**) and the auxin mutants (**F**) taken on day 13. **G.** Changes in regeneration outcomes from day 13 (d13) to day 20 (d20) for each auxinole treatment and auxin mutant. **H.** Example images of a callus-forming explant (left), 1 RR explant (middle), and 2 RRs explant (right).

We began by testing the effect of different concentrations of auxinole on various metrics of root regeneration, such as timing of RR emergence and number of RRs. After removing and culturing the leaf explants, we screened the number of RRs from each explant every day for 13 days. The auxinole treatment delayed the onset of RRs, with the control explants starting to show emerged RRs as early as day 4, while the seedlings treated with 5 μM auxinole began to show emerged RRs on day 6 (Fig. 4B). Regardless of the treatment, once the first RRs emerged, there was a steady increase in the average number of RRs over the 13 days of screening, the slope of which decreased with higher auxinole concentration. To complement the auxinole treatments, we also analyzed the *IAAslow* mutant and a *tir1*^D170E and M473L^ (hereafter referred to as *tir1-D*) double mutant which is hypersensitive to auxin^43^. In the *IAAslow* background, the earliest emerged RRs were delayed similarly to the auxinole treated seedlings (Fig. 4C). In contrast, the *tir1-D* mutant was not delayed and showed a large jump in the average number of RRs per explant between day 5 and day 7 before leveling off to reach a more modest incline from day 8 to day 13 (Fig. 4C). Total number of roots was also correlated to auxin status, with a lower average number of RRs on day 13 for both auxinole treatments and *IAAslow* mutants when compared to control, and a higher average number of roots in *tir1-D* mutants (Fig. 4D). The trend did not reflect viability differences between the conditions, as the trend was similar when analyzing all surviving explants (green) or only explants that regenerated at least 1 RR (yellow).

We next plotted the distributions of number of RRs on day 13 for each explant in auxinole treatment (Fig. 4E). In the mock treated WT, the distribution was the most spread out, with explants regenerating anywhere from 0 to 4 roots at comparable frequencies. With 1 or 2 μM, surviving explants either regenerated 0 roots or they regenerated 2-3 roots. With 5 μM auxinole, over half the explants did not survive to 13 days, and the ones that did regenerated either 0 or 1 roots. Repeating this for the *IAAslow* and *tir1-D* mutants (Fig. 4F), we found the *IAAslow* result to be similar to that of the 5 μM auxinole treatment, where explants either regenerated 0 or 1 roots, but a higher proportion of *IAAslow* explants survived the 13 days compared to the 5 μM auxinole. In the *tir1-D* mutant, somewhat counterintuitively, we found the distribution to be similar to that seen for the 1 and 2 μM auxinole treatments. A higher proportion of *tir1-D* explants survived to 13 days compared to the 1 and 2 μM auxinole treatments, but among the surviving explants, around half regenerated zero roots in all three cases, while the other half regenerated multiple roots (2-3 RRs for 1 and 2 μM auxinole, 2-4 for *tir1-D*).

For every case tested, there were a number of explants that, by day 13, had not regenerated any roots but remained alive, hereafter deemed non-regenerating (NR) explants. Because the NR explants were noticeably lighter in color than the regenerating explants and had a hard, brittle texture (Fig. 4G) matching descriptions of plant callus^44^, we posited that these explants may be on a callus forming trajectory. Callus formation is another means of regeneration where a high auxin signal induces the formation of a pluripotent mass of cells from which regeneration of organs can occur if cultured on appropriate media after callus induction^42^. In lab settings, callus is typically induced by culturing plant tissue in a high auxin media, but callus formation can also be triggered by high auxin after wounding^45^. To more thoroughly assess the regeneration outcomes for the NR explants, we cultured all the explants for an additional week (to day 20). We posited that NR explants committed to a callus trajectory would remain alive without regenerating additional roots by day 20, while NR explants not undergoing callus formation would die off in that time. Because callus formation is associated with very high auxin signaling, we predicted that the *tir1-D* explants, which should have the highest auxin signal, would be most likely to form callus. Indeed, we found that *tir1-D* explants had the highest proportion of NR explants on day 20 (Fig. 4H). Additionally, the proportion did not change between day 13 and day 20, meaning every NR explant on day 13 ended up in the callus forming trajectory. The mock treated WT, representing the second highest auxin signaling condition tested, also did not show a reduction in the proportion of NR explants from day 13 to day 20. In contrast, for every other condition (Fig. 4H), there was a reduction in the proportion of NR explants.

## CONCLUSION

Auxin serves to coordinate cell responses both by rapidly turning on LR initiation genes in founder cells, and by repressing the expression of these genes in neighboring XPP cells^33^. Proper balance between these processes is needed to establish defined boundaries between differentiating and non-differentiating cells, both physically and in terms of cell identities. In the WT auxin conditions, robust establishment of these boundaries enables rapid and coordinated progression from an undifferentiated cell state, through the transition state, eventually reaching a differentiated cell state (Fig. 5, top). Dampening the auxin signal reduces both effects, resulting in a muddying of the cell identities and, therefore, more cells expressing the founder cell marker *GATA23* and increased variability in expression patterns. This dampening, in the case of the *IAAslow* mutant, was associated with increased frequency of *GATA23-*expressing cells which appeared more like XPP cells than LR cells. We posited that this effect was due to stabilized transition state dynamics imbued by the perturbation to auxin signaling, and that these cells represent a transition state in LR development. Overall, in reduced auxin signaling conditions, cells were less likely to differentiate and more likely to take on a transition state identity, potentially due to stabilization of the undifferentiated state and the transition state and destabilization of the differentiated cell state (Fig. 5, middle). Further reduction in the auxin signal all but eliminates establishment of boundaries between differentiating and non-differentiating cells, making differentiation much less likely and much more variable if it does occur (Fig. 5, bottom).

**Figure 5:**
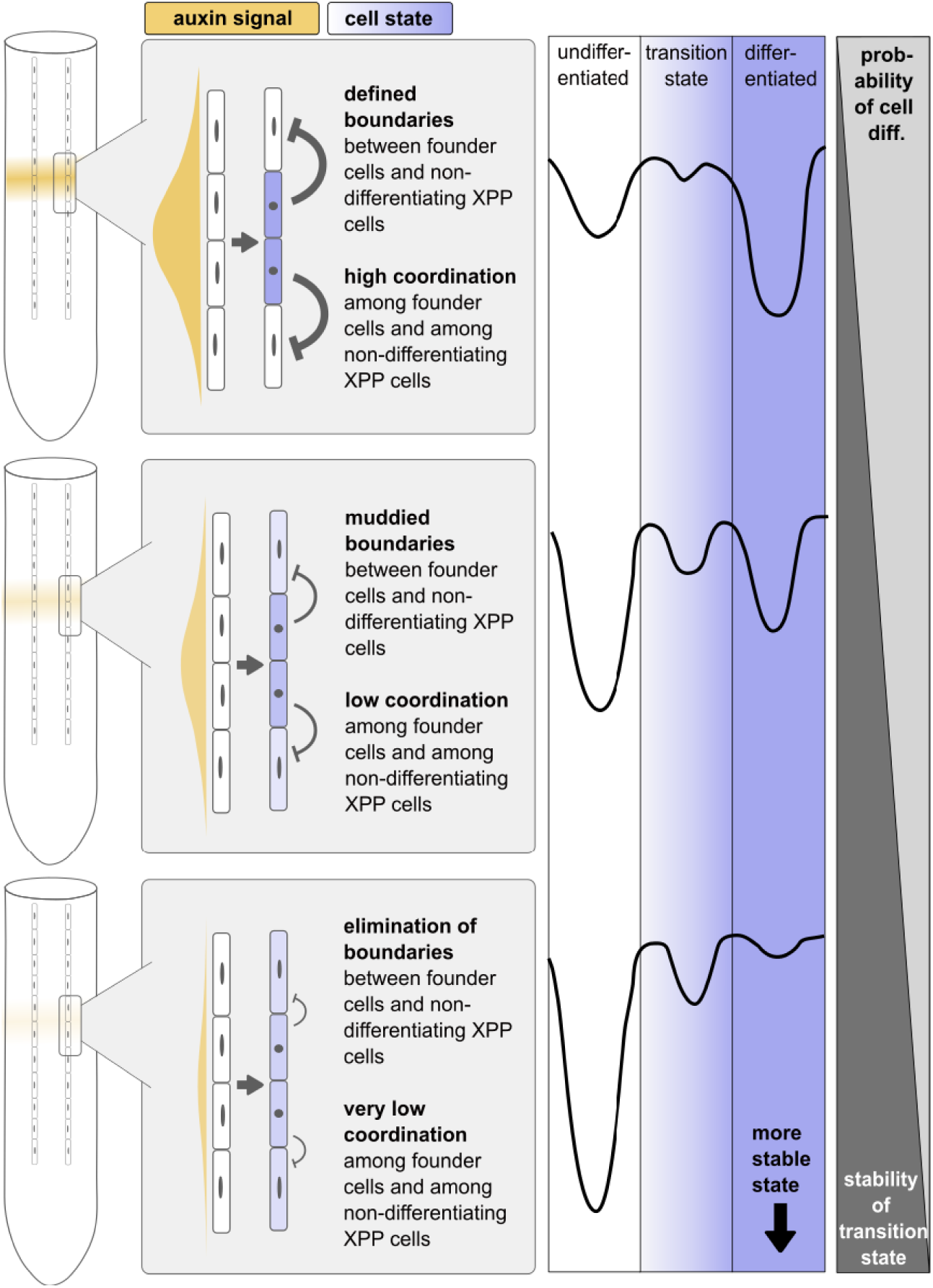
Model of cell differentiation dynamics in different auxin signaling conditions during LR initiation. On the right, graphical representation of the establishment of boundaries between differentiating (purple) and non-differentiating (white) cells with different auxin signal strength (yellow). On the left, representation of energy dynamics underlying undifferentiated, transition, and differentiated states during LR initiation, with deeper valleys representing more stable cell states. Cells are more coordinated when moving through unstable transition states. In a typical auxin signaling context (top) the physical boundaries between XPP cells and LR founder cells is well established through a balance between auxin signal response in founder cells and the repression of this response in neighboring XPP cells. This results in rapid and coordinated progression of founder cells through the transition state to become differentiated. Reduction of the auxin signal (middle) reduces both the strength of the response in founder cells and the repression strength in neighboring XPP cells, resulting in muddied boundaries between differentiating and non-differentiating cells. This reduces the likelihood of differentiation which could be due to stabilization of the undifferentiated and transition cell states and destabilization of the differentiated state. Extreme reduction in the auxin signal (bottom) exacerbates these effects further, resulting in elimination of boundaries and making differentiation even less likely.

Stochastic models of cell differentiation have supported the relationship between signaling conditions and transition state stability and shown that a cell’s differentiation trajectory depends on these dynamics^15–17^. Modeling the response of XPP cells to high auxin concentration^46^ has shown that the increased cellular auxin concentration itself is not sufficient to explain the sustained high auxin response required for these XPP cells to become LR founder cells later during development. Instead, the auxin signal prompts these cells to take on a new “primed” cell state which exhibits a sustained auxin response. This is paired with lateral inhibition mechanisms which prevent nearby LR formation, which is in line with our hypothesis that LR initiation depends both on amplification and attenuation of the auxin response at different sites.

Cell fate decisions in plant cells are quite plastic, with cells along the same lineage undergoing mixed cell fate transitions in certain circumstances^39^. Perhaps a consequence of this plasticity, plant cells also have remarkable regenerative capacity, with differentiated cells under the right conditions becoming totipotent^47^. Auxin promotes cell plasticity, promoting mixed cell fate decisions and regenerative capacity. Work in *Arabidopsis* sepals has shown that cell-to-cell variability in auxin response is needed to initiate development, yet too much initial variability manifests as loss of phenotype robustness, as the variability is too high to be attenuated through typical strategies.

In plants, there is an established connection between hormone signaling dynamics, cell-to-cell variation, cell differentiation, and developmental outcomes, yet engineering efforts typically focus on individual signaling components to achieve desired phenotypes. Additionally, crops are often intentionally engineered with traits that are optimized for a given growth condition, such as a deeper-penetrating root for drought conditions^48^, resulting in low variation in these traits which may not be ideal for growth in increasingly unpredictable environmental conditions due to climate change^49^. Approaches targeting cell-to-cell variation could serve as an alternative approach for engineering developmental traits. Variation could be modulated through changes to the growth environment that alter hormone signaling dynamics and thus may also affect cell-to-cell variation and resilience. Alternatively, genetic modifications to components of hormone signaling pathways could be screened for their effect on cell-to-cell variation, with the goal of identifying lines with increased variation which could confer developmental flexibility or resilience to environmental stressors. This engineering philosophy aligns with Ashby’s Law of Requisite Variety^50^, a foundational principle of cybernetics which states “Only variety destroys variety”. That is, as the complexity of the environment grows, so too must the complexity of the biological response, and engineering cell-to-cell variation could help to achieve this.

## METHODS

### Plant growth conditions

Arabidopsis seeds were sown in 0.5 x Linsmaier and Skoog nutrient medium (LS) (Caisson Laboratories) and 0.8% w/v Phyto agar (PlantMedia/bioWORLD), stratified at 4 °C for 2 days, and grown in constant light at 22 °C.

### Cell counting experiments

Seeds from our cell counting Arabidopsis lines (GATA23 reporter, GATA23 integrase switch34, and GATA23 reporter x IAAslow30) were sown on 0.5X LS Phyto agar plates as described above, one plate with no auxinole added and one plate with 0.5 μM auxinole (MedChemExpress, solid+solvent). After 8 days of growth in constant light at 22 °C, the plates were scanned using a flatbed 767 scanner (Epson America, Long Beach, CA) to assess root length and emerged LR density for each seedling. Seedlings were mounted on slides using diH2O and imaged with a Leica Biosystems DMI 3000 fluorescent microscope (using the RFP channel for the reporter, CFP and RFP channels for the switch, and the GFP channel for the IAAslow reporter cross line). To enable LR width measurements, for the first GATA23 reporter replicate, an image of every LR on each seedling was taken. For all other replicates, all LRs on each seedling were staged as per Banda et al. 2019, but images were only taken of early stage LRs (stage I or II). All LR images were taken at 20X magnification, with the exception of some LRs with enough GECs/GSCs that reduction to 10X was needed to fit them all in view. For a given cell counting line, imaging of both the mock and the auxinole-treated seedlings was done on the same day. Two independent replicates were performed on separate days for each line.

### Regeneration assay

Arabidopsis seeds were sown as described above and grown for 12 days in constant light. The first true leaves from 12 day old seedlings were removed as explants with a doubled edged razor blade (Personna). Explants were then cultured on 0.5X LS plates with 0.8% w/v Phyto agar with concentrations of 0, 1, 2, and 5 μM auxinole. The plates were scanned daily for 13 days of explant culturing using a flatbed 767 scanner and the number of regenerated roots of each explant was also recorded daily. Explants were cultured for an additional week (to day 20) at which point survival, number of regenerated roots, and any callus formation was evaluated and recorded. Callus formation was indicated by a surviving explant without any regenerated roots, with these explants appearing lighter in color and with a brittle texture in comparison to explants with regenerated roots.

### LR/cell counting imaging analysis

All microscope image analysis was performed using the Fiji ImageJ program (version 1.53c). For measuring LR widths the straight line tool was used to span the width of the LR and the measure function was used to measure the distance. For early stage LRs without protrusion, width was assessed by the extent of bright GATA23 reporter expression. For later stage LRs, width was measured from the points of protrusion of the LR from the main root. To assess the number of GECs/GSCs, cells in early stage LRs were counted as GECs if they were sufficiently brighter in reporter expression than any background level expression in the area, and counted as GSCs if they had any mScarlet expression. For stage I LRs, GEC/GSC number was simply the number of expressing cells in the LR. For stage II LRs, having undergone the first periclinal division of LR development, GEC/GSC number was assessed by backtracking to the expected number in stage I, using the well understood patterns of cell divisions in early LR development. For the IAAslow cross line, the shape of the nuclei in each imaged LR was also recorded along with GEC number. LRs containing cells with round nuclei were labeled “LR-like”, while LRs containing cells with oblong nuclei were labeled “XPP-like”. LRs containing cells with both appearances were labeled “mixed”. To assess LR density, seedling plate scans were opened in Fiji ImageJ and root lengths were measured using the segmented line tool.

All plots were generated using Python scripts with plotting functions and was run in version 3.9.1 and with the following package dependencies: pandas (version 1.5.3), scipy.stats (version 1.10.0), matplotlib.pyplot (version 3.6.3), matplotlib.colors (version 3.6.3), and numpy (version 1.24.2).

## DATA AVAILABILITY

Data supporting the findings of this work are available within the paper and its Supplementary Information files. Plasmids and plant materials are available upon request from JLN (; please expect a response within 3 weeks). Source data are provided with this paper.

## ACKNOWLEDGEMENTS

We thank Dr. Alexander Leydon, Janet Solano Sanchez, Benjamin Downing, Julie Ray, as well as other members of the Nemhauser, Imaizumi, Di Stilio, Bagheri, and Steinbrenner groups, for feedback and discussions.

## STUDY FUNDING

Our work on the role of auxin in controlling transcriptional noise is supported by the National Institutes of Health (R35-GM148135) and the Benjamin D. Hall Endowed Chair in Basic Life Sciences.

## CONFLICT OF INTEREST STATEMENT

The authors have no conflicts of interest to declare.

## Supplementary Figures

**Figure S1:**
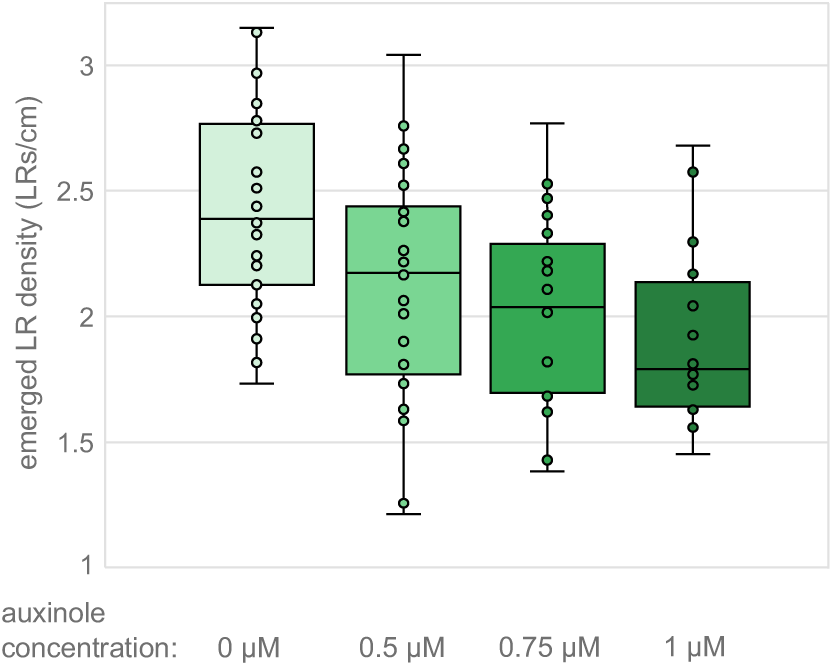
Dose response for emerged LR density in different auxinole concentrations. Seedlings were screened for the number of emerged LRs per cm of main root in 0, 0.5, 0.75, and 1 μM auxinole concentrations. Each point represents the LR density of one seedling in the given auxinole condition.

**Figure S2:**
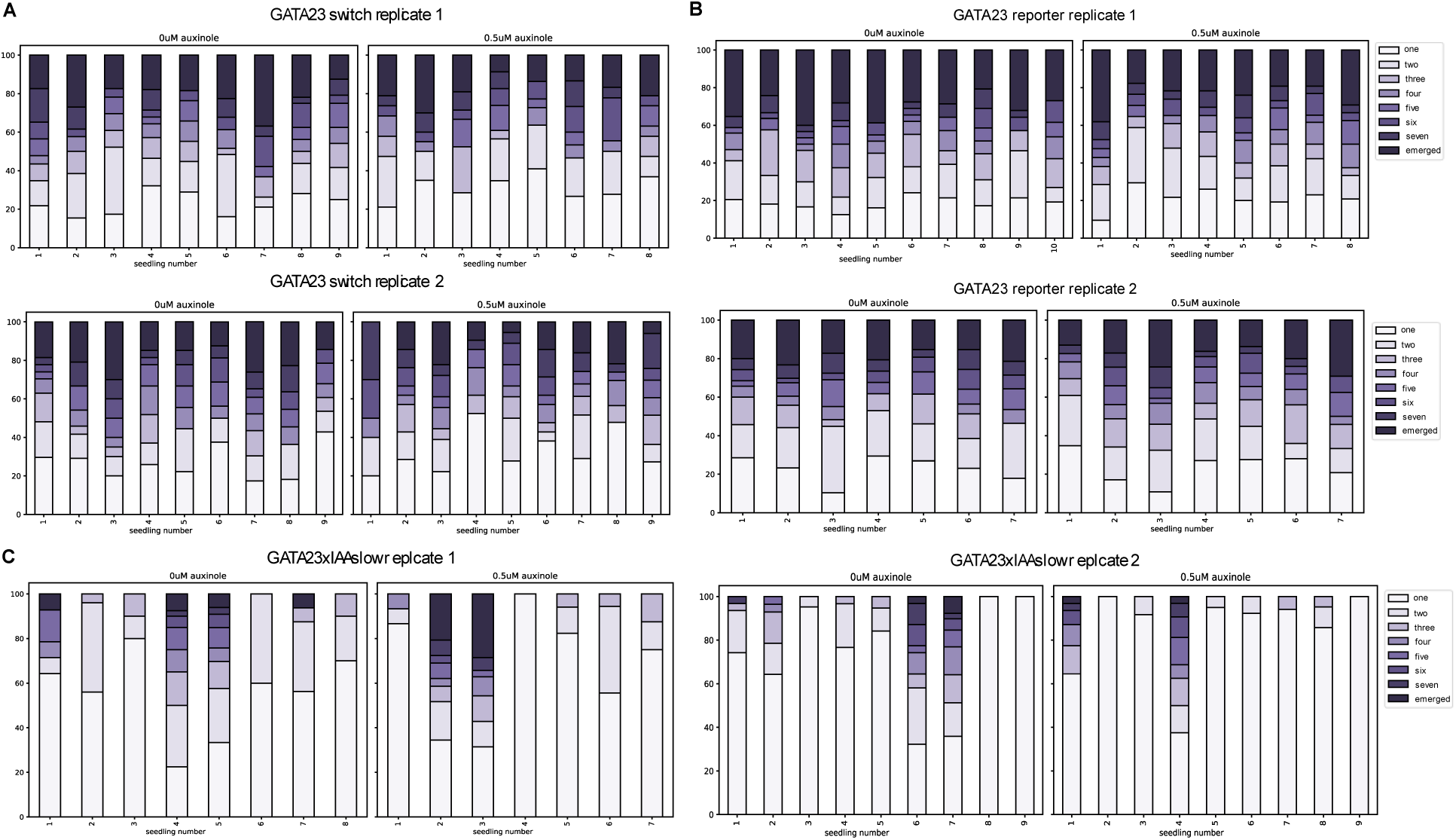
LR staging distributions for *GATA23* switch, *GATA23* reporter, and *GATA23xIAAslow*, auxinole treatment and control. LR staging distributions for each screened *GATA23* switch (A), *GATA23* reporter (B), and *GATA23xIAAslow* (C) seedling across 2 replicates and treated with 0 and 0.5 μM auxinole. Later LR stages are indicated with darker shades of purple as indicated in the legend.

